# Engineering a functional NifDK polyprotein resistant to mitochondrial degradation

**DOI:** 10.1101/755116

**Authors:** Rob Allen, Christina Gregg, Shoko Okada, Amratha Menon, Dawar Hussain, Vanessa Gillespie, Ema Johnston, Andrew Warden, Matthew Taylor, Michelle Colgrave, Keren Byrne, Craig Wood

## Abstract

To engineer Mo dependent nitrogenase function in plants expression of proteins NifD and NifK will be an absolute requirement. Although mitochondria have been established as a suitable eukaryotic environment for biosynthesis of oxygen-sensitive enzymes such as NifH, expression of NifD in this organelle has proven difficult due to cryptic NifD degradation. Here we describe a solution to this problem. Using molecular and proteomic methods, we found NifD degradation to be a consequence of mitochondrial endoprotease activity at a specific motif within NifD. Focusing on this functionally sensitive region, we designed NifD variants comprising between one and three amino acid substitutions and distinguished several that were resistant to degradation when expressed in both plant and yeast mitochondria. Nitrogenase activity assays of these resistant variants in *E. coli* identified a subset that retained function, including a single amino acid (Y100Q) variant. The Y100Q variant also enabled expression of a NifD(Y100Q)-linker-NifK translational polyprotein in plant mitochondria, confirmed by identification of the polyprotein in the soluble fraction of plant extracts. The NifD(Y100Q)-linker-NifK retained function in *E. coli* based nitrogenase assays, demonstrating this polyprotein permits expression of NifD and NifK in a defined stoichiometry supportive of activity. Our results exemplify how protein design can overcome impediments encountered when expressing synthetic proteins in novel environments. Specifically, these findings outline our progress toward the assembly of the catalytic unit of nitrogenase within mitochondria.

## Introduction

Industrial nitrogen fixation is the largest contributor to the world’s population expansion over the past century (1). It is remarkable that this process requiring high temperature (>400°C) and pressure (150 atm) can be accomplished at room temperature by the bacterial enzyme nitrogenase. Hence the notion of harnessing this enzyme for ammonia production in plants has been a longstanding goal of biotechnology (2). More recently, increasing environmental pollution associated with fertilizer production and usage, and food security concerns have lent urgency to this aspiration (3).

Nitrogenase is one of the most complex metalloproteins known in nature (4). The mature enzyme contains a MoFe cofactor at the active site within an α_2_β_2_ heterotetramer comprised of NifD and NifK subunits. The Fe protein, also denoted here as NifH, is required for electron transfer to NifDK. Nitrogenase biosynthesis and function also requires a range of Nif proteins for assembly of the metal clusters and electron transport to NifH. Further to this biosynthetic complexity, both NifDK and NifH are irreversibly destroyed by oxygen (5). These difficulties facing functional reconstitution of nitrogenase in plants have mostly dissuaded attempts towards this goal.

Recently, however, several significant advances in engineering nitrogenase componentry in eukaryotes have been made. Firstly, expression and *ex vivo* function of NifH in yeast mitochondria validated this subcellular location as suitable for reconstitution of oxygen-sensitive nitrogenase components (6). In addition, functional NifB, another oxygen-sensitive enzyme and an essential component of metal cluster biogenesis, has been successfully produced in yeast mitochondria (Buren et al., 2017). Finally, individual expression of 16 Nif proteins from *Klebsiella oxytoca* in plant mitochondria has demonstrated the feasibility of targeting all the biosynthetic and functional Nif proteins to this organelle, although mitochondrial processing and expression was suboptimal for certain members (7).

Despite these advances in producing functional NifH and NifB within mitochondria, expression of NifD remains a problem in both yeast and plants (7, 8). In yeast mitochondria, Burén and colleagues were able to produce NifDK tetramers, but unable to reconstitute nitrogenase function. It was observed that the NifD protein was susceptible to degradation, producing a truncated protein. This degradation product also co-purified with the NifDK tetramer, possibly accounting for the lack of observed NifDK function. Our previous work targeting *K. oxytoca* NifD, to plant mitochondria has also identified this same degradation phenomenon, with the occurrence of a ~48 kDa NifD degradation band as the dominant product. In addition to degradation we were also unable to express NifD in *N. benthamiana* at a similar abundance as NifK. This imbalance presented a further problem as these catalytic partners are required to be expressed in a 1:1 stoichiometric ratio for optimal nitrogenase activity. To address this problem we designed a translational NifD-linker-NifK polyprotein, allowing expression of the polyprotein in a 1:1 stoichiometric ratio. Although function of this polyprotein was untested, a similar translational fusion strategy between NifD and NifK assayed in bacteria retained function (Yang, Xie et al., 2018). This encouraging result indicated that translational fusions of Nif proteins can both simplify the genetic componentry and retain nitrogenase function.

Here we address the issue of NifD degradation within mitochondria. Using molecular and proteomic analyses we discovered that a specific motif within NifD is cleaved by the mitochondrial processing peptidase (MPP) to produce a smaller (~48 kDa) product. We found this site of degradation lies within one of the most functionally sensitive regions within the protein. To address this conundrum, we considered both NifD protein structure and the substrate specificity of the MPP to design variants that were both resistant to degradation and functional. Finally, we show that a translational fusion of NifD and NifK that is resistant to degradation enables expression of a soluble polyprotein in mitochondria, that when assayed in *E. coli,* retains nitrogenase function.

## Results

### NifD is subject to secondary cleavage within plant mitochondria

Previous attempts to express NifD in plant mitochondria produced a ~60 kDa NifD protein of low abundance and a distinct lower molecular weight band of ~48 kDa was observed (7). This ~48 kDa product could be undesirable for several reasons, including interference with formation of the NifDK catalytic unit. We firstly wanted to test whether differing mitochondrial targeting peptides (MTPs) or constitutive promoter combinations may influence the accumulation of this secondary product. For this purpose, we reconfigured our cloning protocols to the GoldenGate system to allow easier interchangeability of these components (9, 10). We then co-infiltrated *N. benthamiana* leaves with various MTP::NifD::HA constructs (Supp. Table 1). Western blot analysis of protein extracts from the infiltrated leaf cells revealed that all constructs produced distinct NifD::HA proteins. However, the ~48 kDa product previously observed was still present, regardless of promoter or MTP, for all NifD expressing constructs.

To determine the origin of the ~48 kDa product we considered two possibilities, downstream translation or protein degradation. The *NifD* gene contains several downstream *ATG* codons which could lead to a ~48 kDa translation product. Therefore, the second to fifth *ATG* codons downstream of the *NifD* start codon were replaced with alternate codons and this protein was expressed as an MTP fusion construct in N. *benthamina* (pRA30, Supp. Table 1). However, the ~48 kDa product was still present in *N. benthamiana* infiltrated protein (Suppl. Fig. 1) refuting the hypothesis that it arose from downstream *ATG* translation. Therefore, we concluded that the ~48 kDa product was degraded NifD protein.

Next, we assessed whether NifD degradation was a consequence of targeting the protein to the mitochondria. To test this, we made two NifD constructs not targeted to the mitochondria, where the MTP sequence was replaced with a HA epitope tag (SN33 and SN34, Fig. 1B). SN34 additionally contained a C-terminal HA tag, allowing detection of any potential C-terminal degradation product. We found that *N. benthamiana* protein extracts from both constructs produced a discrete band of the size expected for full-length NifD proteins (Fig. 1B). The protein band for SN34 was larger than the protein band for SN33 due to the presence of the additional C-terminal HA epitope in SN34. Importantly, there was no ~48 kDa C-terminal degradation product observed after introduction of SN34, in contrast to the mitochondria targeted NifD, SN10. Also there was no N-terminal degradation product observed for SN33 or SN34. Together these results demonstrated that the degradation of MTP::NifD was a consequence of targeting to plant mitochondria.

**Figure 1.**
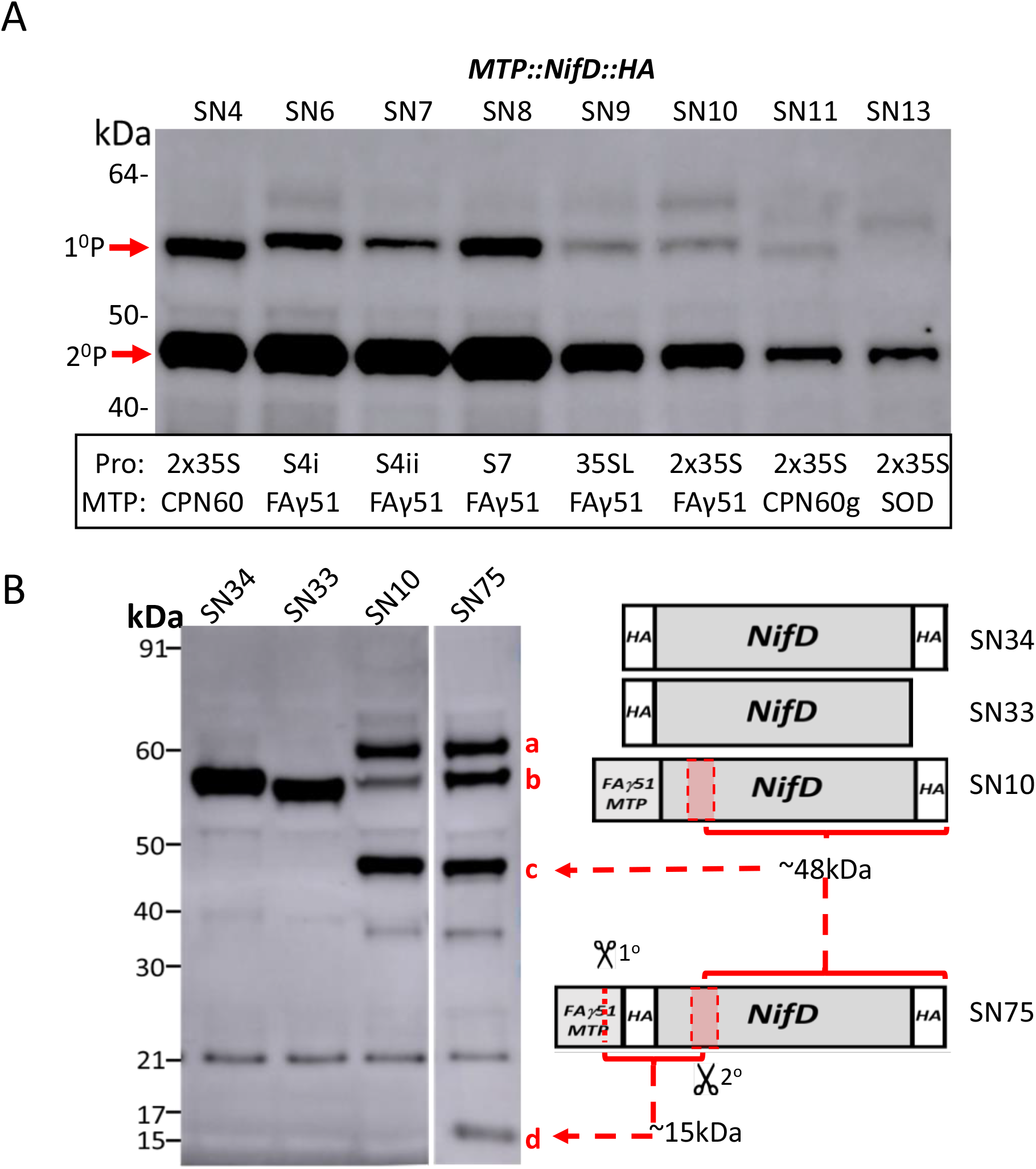
A mitochondrial endoprotease degrades NifD. **A** Western blot (α-HA) of protein extracts after introduction of *MTP::NifD::HA* (SN) constructs into *N. benthamiana* leaf cells. A series of different promoters and MTPs were used to mitochondrially target each SN construct, described in Suppl. Table 1. 1°P = correctly processed MTP::NifD::HA, 2°P = ~48 kDa NifD secondary product. **B** Western blots (α-HA) of protein extracts for mitochondria (SN10, 75) or cytoplasmic targeted NifD (SN33, 34). A schematic of the constructs is shown (not to scale). Red dashed arrows indicate C- and N-terminal cleavage products found for SN75. Red shaded boxes represent an approximate area (not to scale) where cleavage was predicted to occur based on the sizes of the degradation products. MTP::NifD::HA products are designated: a; unprocessed, b; correctly processed, c; secondary C-terminal ~48 kDa cleavage product; d; secondary N-terminal cleavage product at ~13 kDa. The band at ~21 kDa is unspecific background

### Mitochondria targeted NifD cleavage occurs via site specific endoprotease activity

We next attempted to determine whether NifD degradation was a consequence of exo- or endoprotease activity. Given we could detect a NifD C-terminal HA tagged degradation product, detection of a corresponding N-terminal degradation product could confirm endoprotease activity. Accordingly, a construct was introduced into *N. benthamiana* (SN75) identical to SN10 except a HA tag was also included directly after the MTP-FAγ51 and before the NifD coding region (Fig. 1B). Thus, endo-specific cleavage could be expected to produce two HA tagged products - the longer ~48 kDa C-terminal product seen previously in MTP::NifD::HA extracts, and a shorter ~13 kDa N-terminal product. Western blotting analysis of protein extracts revealed a specific shorter ~13 kDa N-terminal product and the longer ~48 kDa C-terminal product (Fig. 1B). This result demonstrated that the cleavage of MTP::NifD was site specific, and not a result of exoprotease degradation from the N-terminus. We therefore describe this ~48 kDa product as a “secondary cleavage product” arising from endopeptidase activity at a region downstream of the MTP processing site.

### The secondary cleavage region within NifD is characteristic of an MPP processing site

The results of the experiments described above indicated that secondary cleavage of MTP::NifD was a consequence of mitochondrial targeting and occurred at a specific location within the NifD sequence. As we could not precisely distinguish this location based on the sizes of the degradation products, we firstly took a broad approach to its identification. For this, a series of MTP::NifD variants were made, each with a block of five consecutive amino acid substitutions (alanine/glycine scanning) within the approximate region (amino acids 49-108) of secondary cleavage within NifD (Fig. 2A). We wanted to test if any of these variants (NifD Var 1-12) would disrupt secondary cleavage.

**Figure 2.**
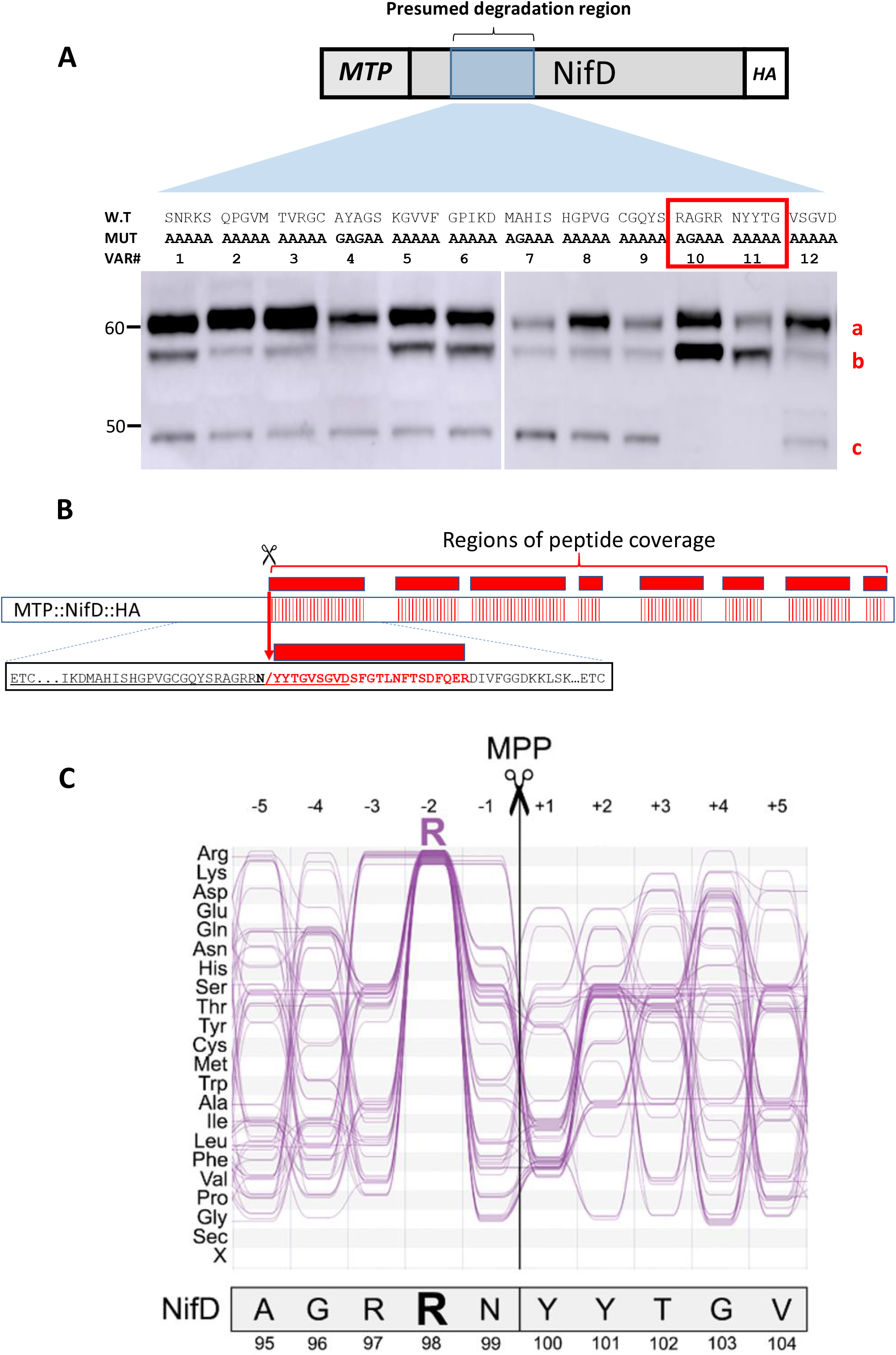
Discovery of the NifD degradation region reveals a characteristic MPP site. **A** Western blots (α-HA) of protein extracts after introduction of MTP::NifD::HA genetic constructs into *N. benthamiana* leaf cells. Constructs were based on SN10 (wild-type NifD), except wild-type amino acids were substituted in five amino acid alanine/glycine blocks (Var 1-12). The initial region for broad coverage with alanine/glycine scanning is shown schematically above (not to scale), with the wildtype NifD sequence, and underneath the corresponding five amino acid change for each individual construct. MTP::NifD::HA bands are designated (a) unprocessed, (b) correctly processed, (c) 2° cleavage product. **B** A schematic of the MS identification of peptides from the degradation product. Eight peptides (red boxes, not to scale) were identified corresponding to NifD sequence. The most N-terminal peptide is shown in detail in red, identifying the precise cleavage site shown by red arrow. **C** Comparison of amino acids 95 to 104 of NifD with a typical −2R MPP cleavage motif. The motif was visualised using selected sequences from *Δicp55* and wild-type *Arabidopsis thaliana,* previously identified by Carrie et al. (19) and Huang et al. (18). For a complete set of mitochondrial cleavage sites, refer to Suppl. Fig. 3. Numbers above refer to the amino acid position relative to the cleavage site. The sequences shown here contain a conserved Arg residue in position −2, characteristic of a −2R MPP cleavage site. NifD also contains an Arg in position −2 relative to the experimentally determined cleavage site.

Of the 12 variants tested, protein extracts for 10 still produced the ~48 kDa secondary cleavage product (Fig. 2A). However, variants 10 and 11 were conspicuous in showing no ~48 kDa cleavage product, and a higher abundance of the correctly processed protein at ~58 kDa. Based on the substitutions in NifD variant 10 and 11, it was concluded that a specific region of the NifD protein within the amino acid sequence RAGRRNYYTG was required for the secondary cleavage of NifD in mitochondria. Using the MitoFates prediction program (11) we found that NifD contained a predicted mitochondrial processing peptidase (MPP) cleavage site in this region, after the asparagine residue, RAGRRN↓YYTG. This motif has similarities to the typical “-2R” motif found in ~55% of mitochondrial targeted proteins (Fig. 2C and Suppl. Fig 3). Therefore, this region of NifD subject to endoprotease activity appeared similar to an MPP cleavage motif.

### Isolation of the NifD degradation product identifies the specific site of endoprotease cleavage

To determine the N-terminal sequence of the degradation product, the ~48 kDa fragment was gel excised and subject to proteomic analysis. In total, eight specific NifD peptides were identified downstream of the region identified from the alanine/glycine scanning approach described above (Fig. 2B). No peptide was found for the complete tryptic peptide RNYYTGVSGVDSFGTLNFTSDFQER, however the expected mass of the shorter peptide YYTGVSGVDSFGTLNFTSDFQER was positively identified. Therefore this shorter peptide must have arisen from cleavage of the NifD sequence within the RRNY sequence – between the asparagine (N) and tyrosine (Y) residues, followed by the tryptic digestion in the analysis. The identification of the cleavage site by this proteomic analysis concurred with the alanine/glycine scanning approach described above and precisely matched the *in-silico* prediction as an MPP cleavage site.

### Design and functional validation of NifD variants resistant to mitochondrial degradation

Although we could abolish secondary cleavage of NifD via five amino acid modifications at the site of MPP activity, it was likely that such changes would disrupt function. We aimed to introduce discrete changes within NifD to potentially reduce or eliminate cleavage whilst maintaining function. Our design was guided by structural modelling and examining the sequence diversity of other naturally occurring NifD sequences at positions corresponding to RAGRRNYYTG of *K. oxytoca*. Our focus was on replacing amino acids in position −3 to +3 relative to the cleavage site (RRN↓YYT). We modelled modifications that would potentially disrupt cleavage while minimising introduction of adverse interactions such as steric clashes, or disruptions to existing hydrogen bonds or electrostatic interactions.

Examination of the crystal structure of the MoFe protein from *Klebsiella pneumonia* revealed that the MPP secondary cleavage site within NifD is in proximity to FeMoco and NifK (Fig. 3A). Specifically, Arg97 interacts with the bridging S5 sulfido ligand of FeMoco and is thought to stabilise the more reduced edge of the cluster (12). Arg98 is involved in a network of hydrogen bonding and electrostatic interactions within NifD and also interacts with Asp516 of the NifK protein. Asn99 accepts a hydrogen bond from Arg97 and Tyr100 interacts with both Arg98 of NifD and Asp516 of NifK, thus contributing to the thermostability of NifDK. We therefore created variants which contained substitutions in the less functionally sensitive residues Tyr101 and Thr102 (i.e. variants 13-15). As we were not confident that these variants described would prevent cleavage by MPP we also included variants that substituted amino acids Arg98 and other sites (Var 27 – 31) recognising these may not be functional.

**Figure 3.**
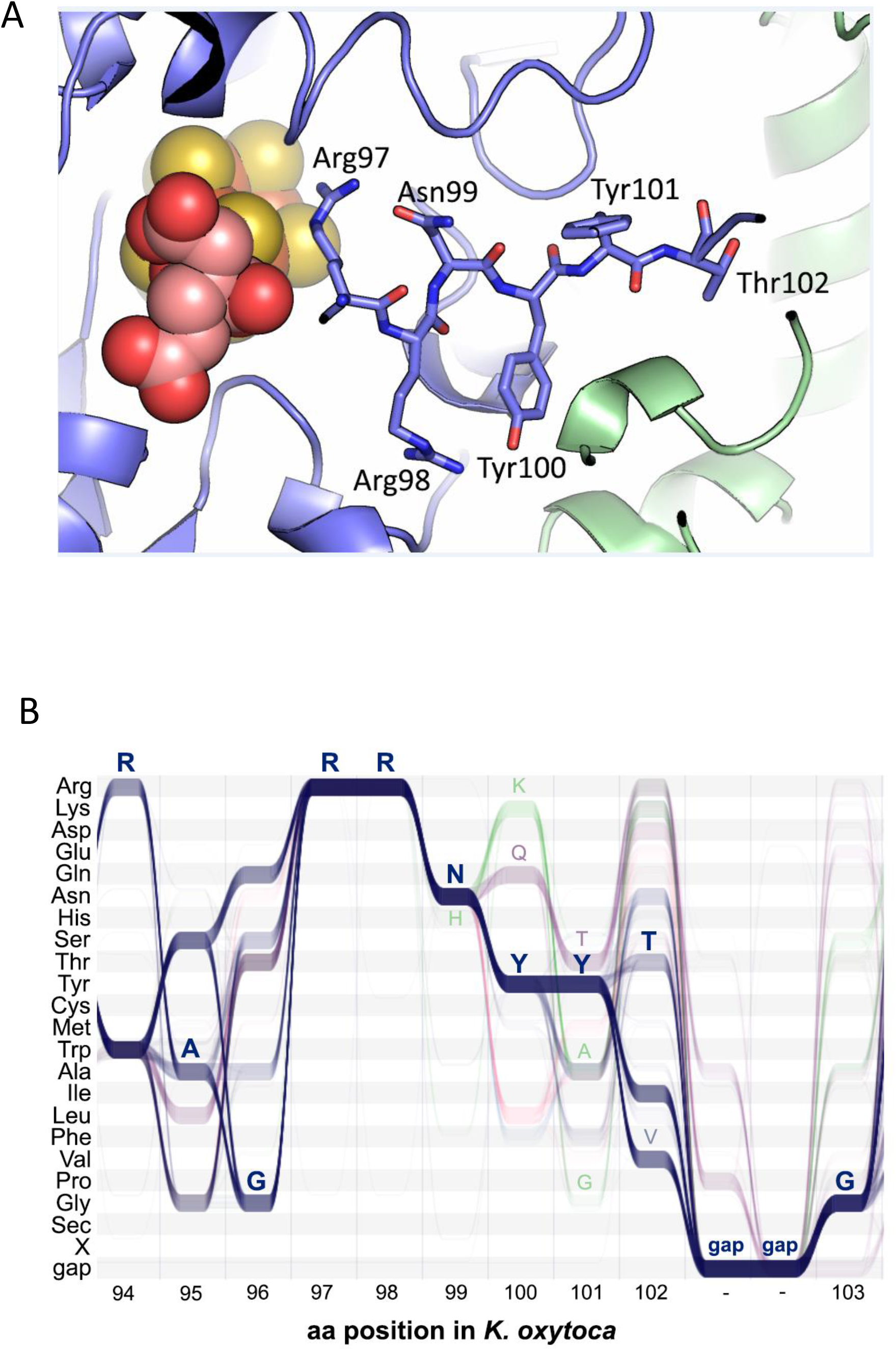
Structural analysis of the RAGRRNYYTG motif in NifD and protein similarity. **A** Location of the proposed secondary cleavage site RRN↓YYT shown in the crystal structure of the MoFe protein from *K. pneumoniae* (PDB:1QGU). NifD is shown in blue and NifK is shown in green. FeMoco is shown as spheres. The residues of the proposed cleavage site are represented by sticks. The cleavage site is in proximity to FeMoco, and to the interface of NifD and NifK. The numbering of amino acids in NifD in the crystal structure was adjusted to correspond to the numbering of the full-length sequence, which contains two methionine residues at the start of the sequence. **B** ALVIS analysis of the amino acid distribution based on 1,476 naturally occurring NifD variants around the proposed secondary cleavage site. The y-axis shows the amino acid residue and the *x*-axis shows the position of the corresponding residues in *K. oxytoca*. The residues of the *K. oxytoca* NifD sequence RAGRRNYYTG are shown above in blue. Sequences that contain a Gln and Lys in position 100 are shown in purple and green, respectively. Amino acid residues that were present in variants are shown above (H99, K100, Q100, T101, A101, G101, V102). Suppl. Table 2 lists the frequency distribution of the equivalent residues of *K. oxytoca* RRNYYT in detail.

Analysis of the sequence diversity among naturally occurring NifD sequences revealed the structural importance of these amino acids was reflected by their conservation (Fig. 3B). For this analysis a set of 1,476 naturally-occurring NifD sequences from a range of bacterial and archaeal sources was extracted from the Interpro database (13) (Suppl. Table 2). The sequences were aligned and visualised using ALVIS (14). The alignment showed that both Arg97 and Arg98 were completely conserved, which is not surprising given their proximity to FeMoco. Overall, the alignment showed that sequence conservation decreases from residue Asn99 to Thr102 (Suppl. Table 2). Accordingly, we also designed variants where these residues were replaced with the second most common residues from the set of NifD sequences, i.e. N99H (Var 18), Y100Q (Var 19), Y101A (Var 26) and T102V (Var 21), respectively. Additionally, amino acids 100 and 101 were replaced with the third and fourth most common residues, i.e. Y100K (Var 24) and Y101T (Var 20), respectively. Furthermore, we noticed some amino acid dependencies and created variants in which two or three amino acids were replaced, i.e. Y100Q/Y101T (Var 22), N99H/Y100K/Y101G (Var 23) and Y100K/Y101A (Var 25).

In total 19 substitution variants were designed (Fig. 4), and these were infiltrated as MTP::NifD::HA (variant) constructs into *N. benthamiana* and leaf protein was analysed by Western blotting. We identified several variants (Fig. 4A) that did not produce the secondary cleavage product (Var 19, 22-25, 29-31). Most remarkably, two variants of this degradation resistant set had single amino acid substitutions at the Y100 residue, Y100Q (Var 19) and Y100K (Var 24). Interestingly some variants (e.g. Var 27) were degraded despite prediction software indicating otherwise (11).

**Figure 4.**
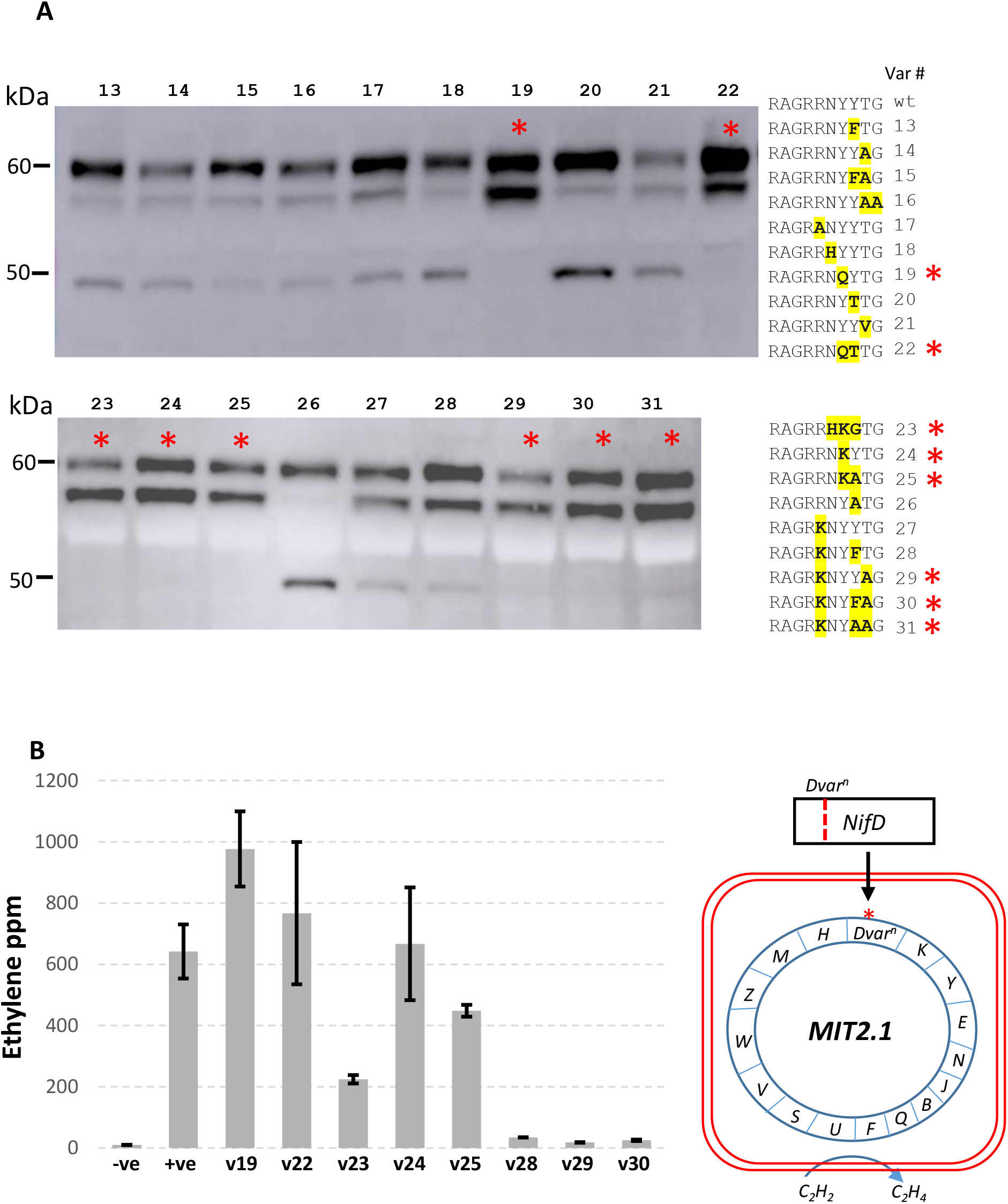
Identification and functional analysis of cleavage resistant NifD variants. **A** Western blot (α-HA) of protein extracts after introduction of *MTP::NifD::HA* variant constructs into *N. benthamiana* leaf cells. Discrete variant sequence changes compared to wild-type are shown in bold/yellow, and variants found to be degradation resistant are identified by a red asterisk. **B** Acetylene reduction assays of nitrogenase activity with variant NifD proteins in *E. coli.* A subset of the same modifications tested above were individually introduced into MIT v2.1 and cultures were grown overnight with 10% acetylene and ethylene was measured. Error bars represent the standard error of the mean from at least two biological replicates. A depiction of the MIT v2.1 assay is shown, where substitutions to NifD are incorporated into the nitrogenase cluster (not to scale, not all genes shown). Negative controls contained no MIT v2.1 cluster, and positive controls included the cluster with the wild-type NifD sequence.

We next wanted to assess the functional consequences of these mutations that provided resistance to NifD degradation. For this purpose, we utilised a nitrogen fixing *E. coli* strain harbouring a modified *K. oxytoca* nitrogenase gene cluster (MIT v2.1) (15) (Fig. 4B). Within MIT v2.1, we introduced mutations in NifD corresponding to a selection of the plant expression variants tested above. We then measured nitrogenase activity of these variants via an acetylene reduction assay (ARA). From our analysis, we identified three variants that retained function similar to wild-type levels (Var 19, Y100Q; Var 22, Y100Q/Y101T; Var 24, Y100K), and others that either had diminished or negligible activity levels (Fig. 4B).

### The Y100Q substitution protects a NifD-linker-NifK polyprotein from mitochondrial degradation at the RRNYYT motif

Our observations that MTP::NifD is subject to secondary cleavage led us to evaluate if degradation is also occurring in a MTP::NifD-linker-K polyprotein (7). For this, we redesigned a NifD-linker-NifK GoldenGate based construct (SN68) differing mainly by placing the HA epitope within the linker peptide avoiding any extension to the C-terminal of NifK. We then introduced the Y100Q variation into SN68 to make the vector SN159. To distinguish processing at the MTP (canonical) cleavage site, a third construct (SN160) similar to SN159 was made except the MTP sequence was modified with alanine substitutions to avoid mitochondrial processing. Finally, an additional control construct (SN176) was designed in which the MTP of SN68 was replaced with a 6xHis tag for expression of cytoplasmic NifD-linker-NifK polyprotein containing the wild-type NifD sequence (Fig. 5B).

**Figure 5.**
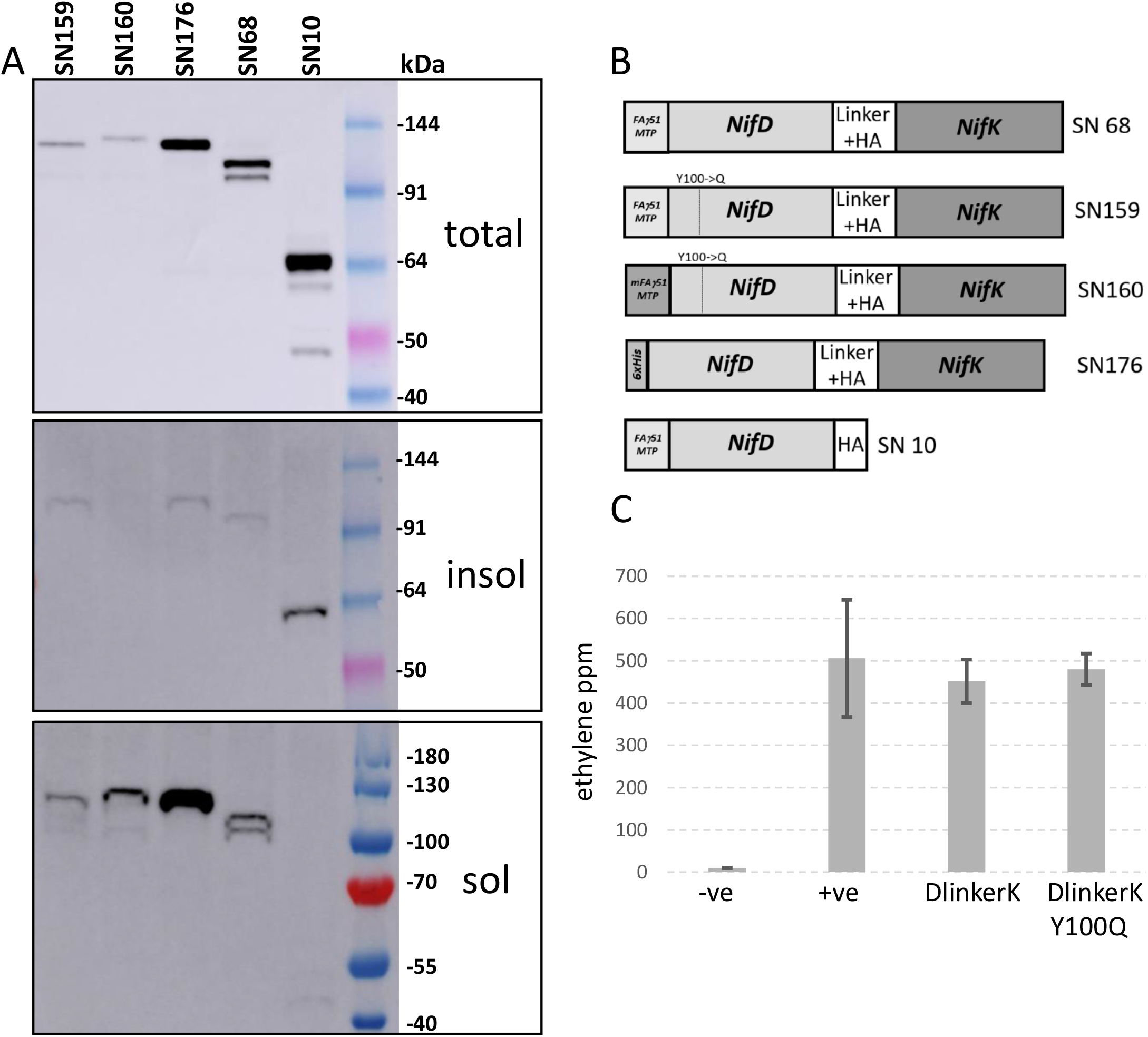
A MTP::NifD-linker-NifK polyprotein is soluble, correctly processed in the plant mitochondrial matrix and functional when expressed in *E.coli.* **A** Western blots (α-HA) of protein extracts after introduction of NifD genetic constructs into *N. benthamiana* leaf cells. Proteins were separated into soluble, insoluble and total fractions (see Material and Methods), a schematic of each construct is described (B). Insoluble and soluble fractions run slightly faster than total fractions due to different buffer conditions. **C** Acetylene Reduction Assays (ARAs) to compare NifD-linker-NifK function to wild-type. For each modification, three biological replicates were measured in duplicate, error bars represent standard error of the mean (SEM).

These constructs were infiltrated separately into *N. benthamiana* leaves and protein fractions were analysed by Western blot. Total protein extracts of the SN159 construct produced a single band that aligned with the SN176 cytoplasmic control, and migrated faster than the SN160 (unprocessed control) (Fig. 5A). This migration speed for the SN159 band corresponded to correctly processed and undegraded MTP::NifD(Y100Q)-linker-K polyprotein. By contrast, the SN68 construct produced two lower bands indicative of degradation. Interestingly, whilst MTP::NifD::HA (SN10) was incompletely processed, the MTP::NifD-linker-K polyprotein was completely processed at the canonical MPP processing site within the MTP (Fig. 5A).

### The NifD-linker-NifK polyprotein is soluble in plant mitochondria and functional in *E. coli*

Solubility of nitrogenase proteins within mitochondria is considered a prerequisite for function. However, a recent report has indicated that targeting *Azotobacter vinelandii* NifB to the mitochondria produced an insoluble protein (16). Therefore, we assessed the solubility of our mitochondria targeted NifD proteins. After preparing soluble and insoluble NifD protein fractions from the same leaf spots used for total protein (see Materials and Methods), we visualised the proportions of proteins in these fractions by Western blot. Whereas a cytoplasmic targeted version of NifD (SN33) was mostly soluble (Suppl. Fig. 2), we were unable to detect MTP::NifD (SN10) in the soluble fraction (Fig. 5A). In contrast, we were surprised to discover that more of the NifD(Y100Q)-linker-NifK polyprotein produced from SN159 was observed in the soluble fraction than in the insoluble fraction. This was also the case for the other NifD-linker-NifK polyproteins, including the cytoplasmic targeted version, SN176. Therefore, fortuitously, the polyprotein from SN159 resistant to the secondary cleavage in mitochondria was also predominantly soluble.

We assessed the function of the NifD-linker-NifK polyprotein by introducing the same linker as used for plant expression (SN159) between the native NifD and NifK proteins from MIT v2.1 and tested for nitrogenase function via ARA. We also introduced the Y100Q variation into this NifD-linker-NifK polyprotein in MIT v2.1. We found that both the NifD-linker-NifK and degradation resistant NifD(Y100Q)-linker-NifK proteins retained wild-type levels of activity (Fig. 5C). These results also concurred with our prior assays of the Y100Q mutation within NifD, demonstrating this mutation within NifD, either when expressed singly, or fused with NifK does not impede nitrogenase function.

### The Y100Q variation within NifD prevents secondary cleavage in yeast mitochondria

Yeast have proven a useful model for testing expression and assembly of nitrogenase components in mitochondria (6, 8, 17), however, the problem of NifD degradation has also been reported when expressing *A. vinelandii* NifD in this organelle. Notably, *A. vinelandii* NifD contains the same RRNYYT motif that enables MPP cleavage of *K. oxytoca* NifD in plant mitochondria. We wanted to determine whether *K. oxytoca* NifD also was similarly degraded when expressed in yeast mitochondria, and secondly whether the Y100Q variation could protect NifD from any possible degradation. For this we made yeast expression vectors targeting *K. oxytoca* NifD to either the cytoplasm (SNY196) or the mitochondrial matrix (SNY10). We also made a yeast expressed version of NifD Y100Q variation (SNY114) (Fig. 6). Besides being expressed via the GAL4 promoter/terminator, these constructs were otherwise identical to their plant equivalents.

**Figure 6.**
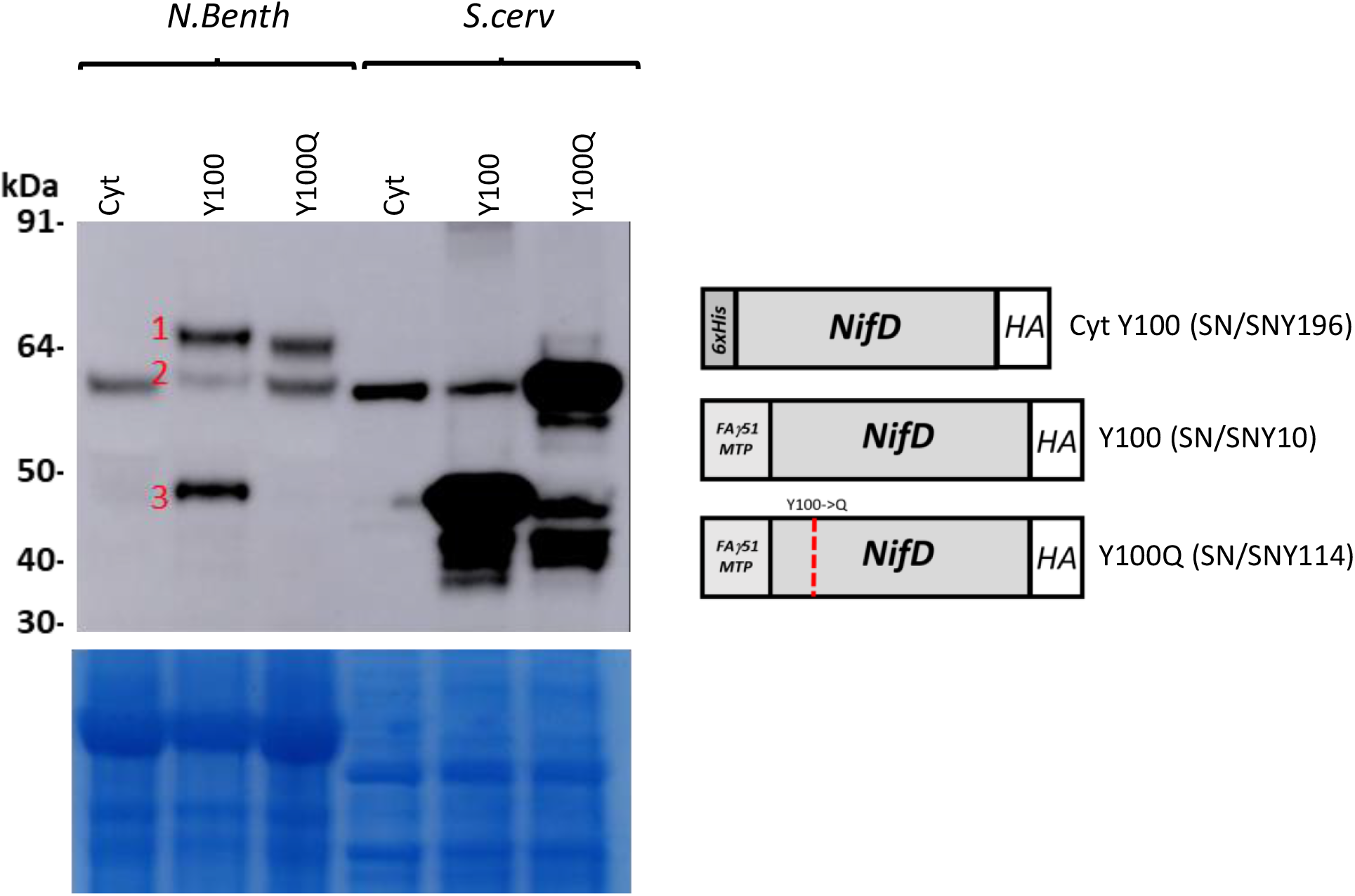
The Y100Q mutation in NifD prevents secondary cleavage in yeast mitochondria. **A** Western blot (α-HA) of protein extracts after introduction of MTP::NifD::HA genetic constructs into *N. benthamiana* leaf cells or *S. cerevisiae.* A schematic of the constructs is shown (not to scale), and the Y100Q synthetic NifD construct resistant to production of the secondary cleavage product is indicated. “Cyt” = cytoplasmic targeted construct. 1= unprocessed NifD band, 2= processed NifD occurring by primary cleavage of the MTP, 3= unprocessed degradation product resulting from secondary cleavage. Coomassie stained gel post transfer shown underneath.

The yeast constructs were expressed in *Saccharomyces cerevisiae*, and total protein was analysed via Western blot. We found both plant and yeast expressing *K. oxytoca* NifD cytoplasmic constructs produced a band of the size expected for cytoplasmic expression of full-length NifD, and no C-terminal degradation products were observed (Fig. 6). However, the yeast mitochondria targeted NifD construct SNY10 produced an intense smaller band of approximately the same size as the plant ~48 kDa degradation product from SN10, whereas much less full length NifD was produced by this construct. This result indicated that *K. oxytoca* NifD degradation was occurring in yeast mitochondria, possibly at the same RRNYYT motif as occurs in plant mitochondria. Notably the NifD Y100Q mutation within yeast mitochondria enabled a much higher level of undegraded NifD to accumulate (Fig. 6), although smaller products were also observed for all proteins targeted to yeast mitochondria. Together these results indicated that the wild-type NifD protein from *K. oxytoca* is degraded in the same region when targeted to yeast or plant mitochondria, and this could be greatly reduced via the Y100Q variation, enabling a higher level of full length NifD.

## Discussion

Mitochondria have potential as an organelle suitable for nitrogenase reconstitution (6), however both NifD instability and insolubility within mitochondria present functional bottlenecks. Our modifications to NifDK have produced a soluble protein that is resistant to secondary cleavage within mitochondria and retains nitrogenase function. This study demonstrates that obstacles facing the transfer of ancient conserved pathways to novel environments can be overcome by approaches such as those taken here.

Several strands of evidence imply the secondary cleavage of NifD is due to the activity of the MPP. Primarily, the identified secondary cleavage site contains the hallmarks of a canonical MPP processing site, namely the presence of an arginine at position −2 in relation to the scissile bond, characteristic of cleavage motifs found in the majority of matrix-targeted proteins (18). Secondly, disruption of this site by alanine/glycine scanning such that the region no longer resembles a canonical MPP cleavage site prevents the secondary cleavage of NifD. Moreover, targeting of NifD to the cytoplasm, which does not contain an MPP, either in yeast and plants, prevents secondary cleavage.

We designed several discrete variants at the identified secondary cleavage site (e.g. Y100Q), that prevented NifD degradation and retained function. However some of these modifications would not have been predicted to avoid MPP activity (11). Conversely, other variants that would be predicted to provide MPP resistance did not (e.g. Var 27). Our results show that although *in silico* predictions are a useful guide, these are ultimately constrained by the fact there is no absolute consensus for an MPP cleavage site (19). This has implications for the future design of synthetic Nif proteins targeted to organelles. Consideration needs to be given to potential MPP sites, but given our ability to predict these is limited, the potential exists for unintended cleavage of synthetic proteins.

Although we could protect NifD from degradation via the Y100Q substitution, this protein was only partially processed at the canonical MPP cleavage site within the MTP. However, the addition of NifK to the C-terminus of NifD enabled complete processing of the polyprotein for unknown reasons. That processing is enhanced by adding the NifK protein sequence to NifD is remarkable, particularly as this synthetic polyprotein (120 kDa) is longer than any known endogenous protein that is transported to plant mitochondria (18). Even larger Nif polyproteins have been shown to function in *E. coli,* and may provide a means to simplify expression and provide correct stoichiometry (20). It is therefore encouraging that mitochondria appear capable of importing and processing larger synthetic proteins, and it will be interesting to test the upper size limits for other nitrogenase polyproteins.

We found that *K. oxytoca* NifD is insoluble when targeted to plant mitochondria, a result similar to that reported for *A. vinelandii* NifB (16). However, the translational fusion of NifD to NifK led to a soluble polyprotein within the mitochondria. It is possible that linking these proteins has allowed rapid association of the NifD and NifK components into their native heterotetrametric structure within the mitochondrial matrix. Furthermore, we show that individually expressed NifD was soluble in the cytoplasm but not in the mitochondria. This observation suggests that the mitochondrial targeting of NifD or the mitochondrial environment itself prevents NifD solubility. As many possibilities exist for why Nif proteins are insoluble when targeted to mitochondria, approaches such as taken by Burén and colleagues (16) to screen naturally occurring variants to identify soluble versions, or here with linking partner proteins, may offer generic solutions to this issue.

We identified that degradation of *K. oxytocoa* NifD occurs in yeast mitochondria, which was reduced by incorporating the Y100Q variation. As *A. vinelandii* NifD also contains an identical internal RRNYYT site, it is likely the degradation seen for *A. vinelandii* NifD when expressed in yeast (8) is a consequence of secondary cleavage at this site. Therefore, we would expect an *A. vinelandii* NifD containing the Y100Q mutation targeted to yeast mitochondria would also show a reduction or elimination of the degradation product. As *A. vinelandii* NifD degradation has been suggested to present a possible bottleneck to formation of the correct NifDK heterotetramer, it will be of interest to see if a degradation resistant version of *A. vinelandii* NifD may enable formation of active NifDK tetramers in yeast when co-expressed with the other cohort of required Nif proteins.

Although nitrogenase biosynthesis and function is complex, significant progress has been made towards reconstructing nitrogenase components within mitochondria. The demonstration that active NifH and NifB can be isolated from yeast mitochondria augurs well for both electron donation and maturation of the metalloclusters within mitochondria. Here we have overcome both degradation and insolubility bottlenecks to express the catalytic unit of nitrogenase within this organelle. Future investigations can now begin to combine these maturation, electron donation and catalytic pathways to explore the function of nitrogenase within mitochondria.

**Supplementary Figure 1.**
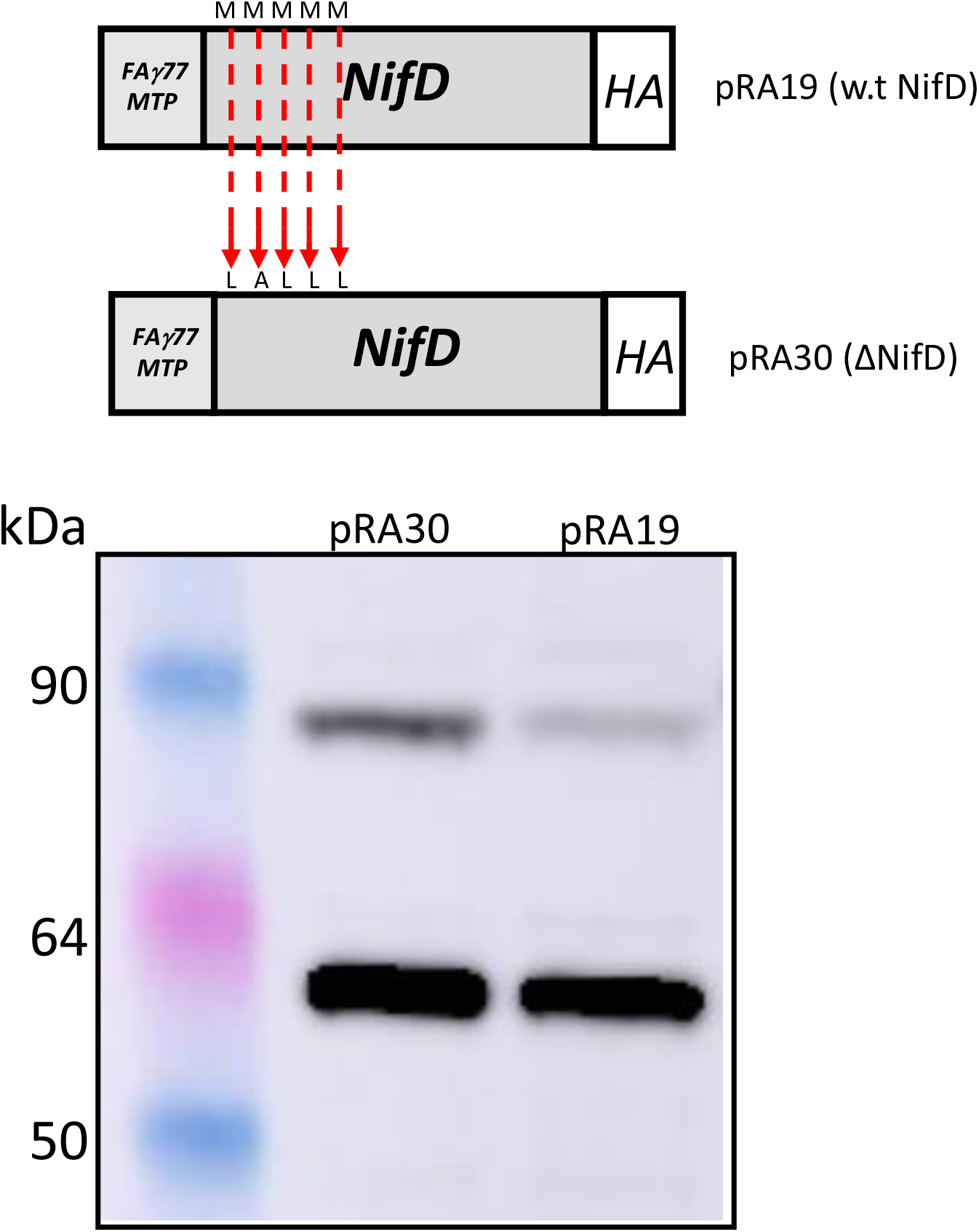
Removal of downstream NifD *ATG* codons does not prevent accumulation of a degradation product. A construct based on pRA19 (7) was modified by replacing the five methionine codons downstream of the start codon with alternate codons (coding amino acids L, A, L, L, L) to create pRA30. After infiltration of this construct and pRA19 separately into *N. benthamiana,* protein extractions and western blotting were carried out using α-HA.

**Supplementary Figure 2.**
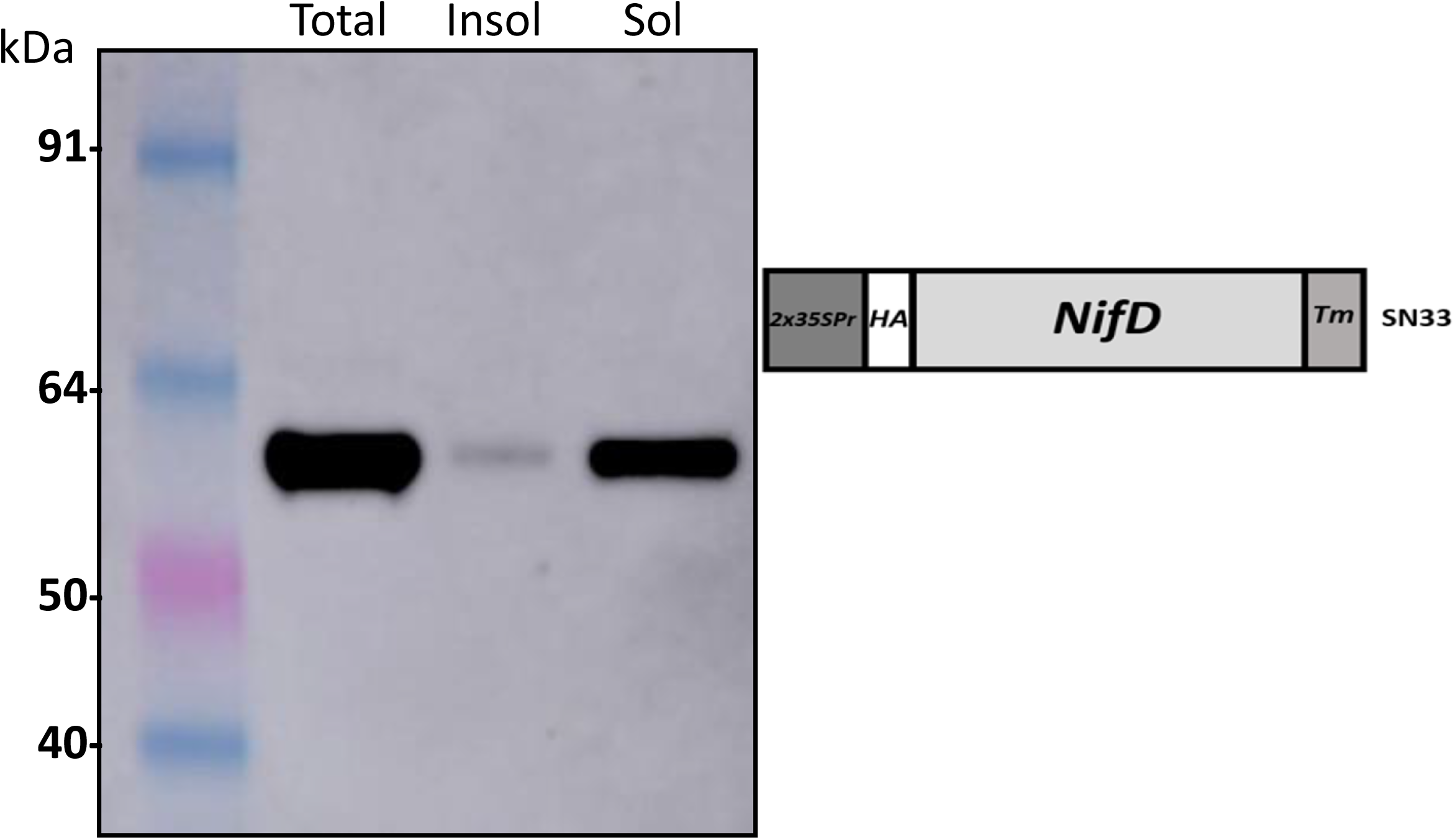
Cytoplasmic targeted NifD accumulates mostly in the soluble fraction. Western blot of cytoplasmic NifD construct. After *N. benthamiana* infiltration, protein was separated in total, soluble and insoluble fractions, and subject to western blot analysis using α-HA. No degradation product was observed and most NifD accumulated in the soluble fraction.

**Supplementary Figure 3.**
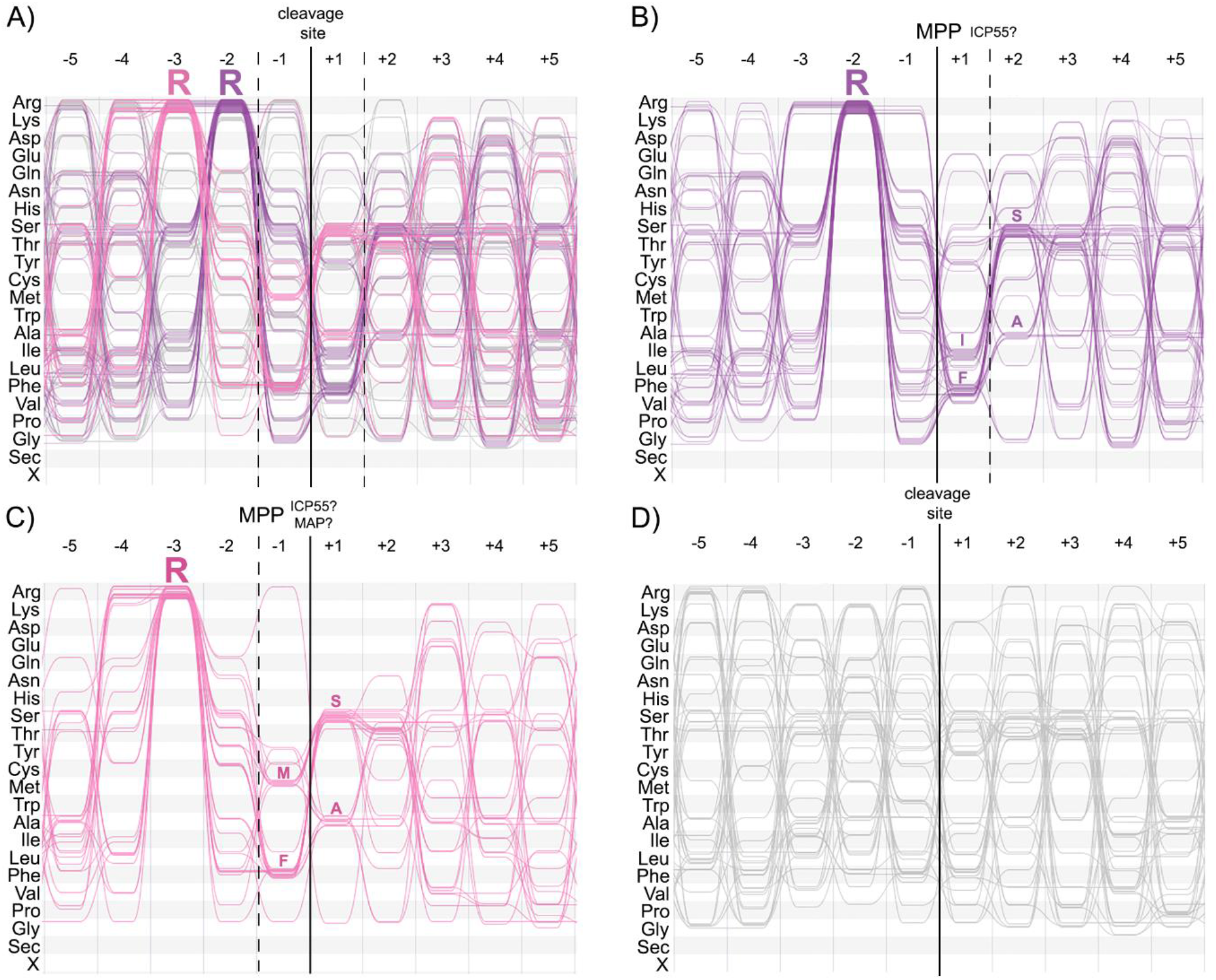
Visualisation of plant mitochondrial cleavage sites. **A** ALVIS visualisation of mitochondrial cleavage sites from *Δicp55* and wild-type *Arabidopsis thaliana,* previously identified by Carrie et al. (19) and Huang et al. (18). Sequences containing an Arg residue in position −2 or −3 are shown in purple and pink, respectively. Sequences containing no conserved Arg residue are shown in grey. Of the 140 sequences in this alignment, 65 belong to the −2R group, 30 to the −3R group, and 45 contain no conserved Arg residue. **B** Alignment of −2R proteins, which are cleaved by the MPP at the indicated site. As some cleavage sites were identified in an Δicp55 background, some proteins contain an additional ICP55 cleavage site between position +1 and +2. **C** Alignment of −3R proteins. The MPP cleavage site is most likely located between position −1 and −2. Cleavage by the MPP is then followed by the removal of one further amino acid. ICP55 has been shown to remove Phe, Tyr and Leu residues (Carrie et al., 2009) that make up 50%, 6.6% and 3.3%, respectively, of the −3R proteins shown here. Met residues (26.6%) could potentially be removed by a methionine aminopeptidase (MAP) (Giglione et al., 2001). **D** Sequences containing no conserved Arg residue in position −2 or −3.

## Materials and Methods

### Construction of Vectors

Vectors for *N. benthamiana* expression were based on the GoldenGate parts and tool kit (10) and assembled using the GoldenGate assembly protocol (9). Non-standard components and NifD variant sequences for plant expression vectors were generated by DNA synthesis. For yeast expression constructs NifD flanking sequences were amplified from SN10, SN114 and SN196 using primers containing KpnI (Forward) and SacI (Reverse) restriction sites for cloning into the yeast expression vector pYES2. For cloning of NifD variant sequences within MIT v2.1, silent mutations that created AgeI and SalI sites were introduced into the NifD coding sequence to create MIT v2.1mod. Then corresponding AgeI/Sal fragments were excised from variant NifD sequences and ligated into the AgeI/SalI sites within MIT v2.1mod. For generation of the NifD-linker-NifK within MIT v2.1 ligase cycling reaction (LCR) was used to fuse the same linker as used in SN68 between the NifD and NifK subunits. Mutagenesis was used to introduce the Y100Q mutation within this vector.

### Plant growth and transient transformation of *N.benthamiana*

*N. benthamaina* plants were grown in a Conviron growth chamber at 23°C under a 16:8 hr light:dark cycle with 90 μmol/min light intensity provided by cool white fluorescent lamps. *A. tumefaciens* strain GV3101 (SN vectors) or AGLI (P19 vector) cells were grown to stationary phase at 28°C in LB broth supplemented with 50 mg/ml carbenicillin or 50 mg/L kanamycin, according to the selectable marker gene on the vector, and 50 mg/L rifampicin. Acetosyringone was added to the culture to a final concentration of 100 μM and the culture then incubated at 28°C with shaking for another 2.5 hr. The bacteria were then pelleted by centrifugation at 5000 x g for 10 min at room temperature. The supernatant was discarded, and the pellet was resuspended in a solution containing 10 mM MES pH 5.7, 10 mM MgCl_2_ and 100 μM acetosyringone after which the OD_600_ was measured. A volume of each culture, including the culture containing the viral suppressor construct 35S:P19, required to reach a final concentration of OD_600_ = 0.10 was added to a fresh tube. For plant transformations with MTP::NifD constructs, MTP::NifK (SN46) was added to the infiltration mix. The final volume was made up with the infiltration buffer. Leaves of five-week-old plants were then infiltrated with the culture mixture and the plants were grown for five days after infiltration before leaf discs were recovered for analysis.

### Yeast transformation

Transformation of yeast strain INVSc1 (Thermo Fisher Scientific) was performed using the Yeast Transformation Kit (Sigma Aldrich) according to the manufacturer’s protocol. Transformed colonies were selected by plating the transformation mixture onto minimal medium without uracil (SCMM-U) agar plates, which contained 6.7 g/L yeast nitrogen base, 1.92 g/L synthetic dropout medium without uracil (Sigma Aldrich), 20 g/L glucose, and 20 g/L agar. After 2-3 days of incubation at 30°C, single colonies were restreaked onto fresh SCMM-U agar plates. A single colony that contained the genetic construct was inoculated into SCMM-U liquid media (containing the same components as SCMM-U agar but without the agar), grown at 30°C with shaking for 2 days. Glycerol was added to a final concentration of 20% and aliquots stored in −80°C until further use.

For expression of the genes contained in the genetic construct, an inoculant from the glycerol stock was grown in SCMMM-U liquid media at 30°C with shaking for 2 days. The cells were collected from the culture by centrifugation and resuspended in SCMM-U induction medium which was identical to SCMM-U liquid media except that the glucose was replaced with 20 g galactose, to a final OD_600_ of 0.4. The culture for induction was grown at 30°C with shaking for 2 days and the yeast cells were collected by centrifugation for protein extraction and Western blot analysis.

### Plant and yeast protein extractions and western blot analysis

For analysis of total protein, extractions were as described in (7). Yeast protein was extracted using the same buffer as for *N. benthamiana* except ~ 100 mg of cells were used with an equivalent volume of extraction buffer. For preparation of soluble and insoluble fractions, from the same infiltrated leaf disc used for total protein, approximately 180 mm^2^ of leaf disc was ground under liquid nitrogen using a mortar and pestle and 300 μL of cold solubility buffer was added. The solubility buffer contained 50 mM Tris-HCl pH 8.0, 75 mM NaCl, 100 mM mannitol, 2 mM DTT, 0.5% (w/v) polyvinylpyrrolidone (average mol wt 40,000), 5% (v/v) glycerol, 0.2 mM PMSF, 10 μM leupeptin and 0.5% (v/v) Tween^®^ 20. The samples were centrifuged for 5 min at 16,000 x g at 4°C. The supernatant was transferred to a fresh tube and the pellet was resuspended in 300 μL of cold solubility buffer. Both, the supernatant (sample 1) and the resuspended pellet (sample 2) were centrifuged again for 5 min at 16,000 x *g* at 4°C. From sample 1, a sample was taken from the supernatant, which is referred to as the soluble fraction. This sample was mixed with an equivalent amount of 2 x SDS buffer. 2 x SDS buffer contained 250 mM Tris-HCl pH 6.8, 8% (w/v) SDS, 40% (v/v) glycerol, 120 mM DTT and 0.004% (w/v) bromophenol blue. After the second centrifugation step, the supernatant of sample 2 was discarded. The pellet is referred to as the insoluble fraction. The pellet was resuspended in 300 μL 4 x SDS buffer and 300 μL of solubility buffer were added. Samples for the total, insoluble and soluble fractions were heated at 95°C for 3 min and then centrifuged at 12000 x *g* for 2 min. 20 μL of the supernatant containing the extracted polypeptides was loaded on a NuPAGE Bis Tris 4-12% gels (Thermo Fisher Scientific) for gel electrophoresis and Western blot analysis, as described by (7).

### Sequence alignment and visualisation of NifD sequences

1,751 NifD sequences (family IPR005972, nitrogenase molybdenum-iron protein alpha chain) were extracted on 12 December 2018 from the InterPro database (13). Removal of duplicate sequences resulted in a set of 1,476 unique sequences. The sequences were aligned using the multiple sequence alignment program MAFFT (version 7) (21). The FFT-NS-2 (fast but rough) strategy was used with default parameters. The aligned sequences were visualised using ALVIS (interactive non-aggregative visualization and explorative analysis of multiple sequence alignments) (14).

### Acetylene Reduction Assays using the pMIT2.1 system in *E. coli*

Cells of *E. coli* strain JM109 were transformed with the plasmids pMIT v2.1 (or one of its derivatives that was being tested) and pN249 which conferred resistance to the antibiotics chloramphenicol and spectinomycin, respectively, as described in (22). The transformed cells were selected by growth on LB medium (10 g/L tryptone, 5 g/L yeast extract, 10 g/L NaCl) containing chloramphenicol (34 mg/L) and spectinomycin (80 mg/L). Transformed cells were grown aerobically overnight at 37°C in LB medium with antibiotics to an optical density of 1. 0. The cultures were centrifuged at 10,000 x g for 1 minute and the supernatant discarded. The cells were re-suspended in one volume of an induction medium which was free of N-sources, containing 25 g/L Na_2_HPO_4_, 3 g/L KH_2_PO_4_, 0.25 g/L MgSO_4_.7H_2_O, 1 g/L NaCl, 0.1 g/L CaCl_2_.2H_2_O, 2.9 mg/L FeCl_3_, 0.25 mg/L Na_2_MoO_4_.2H_2_O and 20 g/L sucrose (minimal medium) supplemented with 1.5 ml/L of 10% serine, 600 μl/L 0.5% Casamino acids, 5 mg/L biotin and 10 mg/L para-aminobenzoic acid (20). The medium was sparged with argon gas for 20 minutes prior to mixture with the bacteria and antibiotics. Stock solutions were filter sterilized. For induction of *nif* gene expression, the medium was supplemented with isopropyl-β-D-1-thiogalactopyranoside (IPTG; Gold Bio#I2481C25 259) at a final concentration of 0. 1 mM, 0.5 mM or 1.0 mM unless otherwise stated, generally 1.0 mM. The cell suspensions were transferred to 3.5 cm^3^ culture flasks and capped with gas-tight rubber seals using a crimp-lock system and the headspace was sparged with pure argon gas for 20 min. The suspensions were then incubated at 30°C with shaking at 200 rpm for 5 hours. After this, acetylene reduction assays (ARA) were started by the injection of 0.5 cm^3^ of pure C_2_H_2_ (BOC gases, instrument grade; final concentration 10% C_2_H_2_ in argon) and further incubation for 18 hours. Production of ethylene at the final time was measured by gas chromatography with Flame Ionisation Detection (GC-FID) using an Agilent 6890N GC instrument. Headspace samples (0.5 cm^3^) were removed and manually injected into a split/splitless inlet on a 10:1 split mode. The instrument was operated under the following parameters: inlet and FID temperatures of 200°C, average velocity for the carrier He of 35 cm/sec, isothermal oven temperature at 120°C. A RT-Alumina Bond/MAPD column (30 m x 0.32 mm x 5 μm) was used with a 5 m particle trap column coupled to the detector end. Analytical performance of the instrument was assessed by running suitable blanks and standards. Under these conditions, ethylene emitted from the column at about 2.3 minutes and acetylene at about 3.1 minutes.

### MS identification of the NifD cleavage product

Protein extracts from *N. benthamiana* leaves infiltrated with SN14 (MTP-Su9::NifD::HA) were run on SDS-PAGE using a gel having a polyacrylamide concentration of 4-20 % (Invitrogen). The gel was stained with Aqua stain (Bulldog Bio). After destaining in water, 5 slices were cut from the gel for the region spanning the molecular weights 37-50 kDa. The slices were numbered 1 to 5 from the smaller molecular weights to the larger. Each gel slice was cut into approximately 1 mm^3^ cubes and soaked in 150 μl 30% methanol for 15 minutes. To reduce proteins that may have oxidised, the buffer was removed and replaced with 100 μl of fresh 25 mM ammonium bicarbonate (ABC) buffer with 5 μl of 15% dithiothreitol and incubated at room temperature for an hour. Cysteine residues were inactivated by the addition of 5 μl of 40% acrylamide and incubation at room temperature for 1 hour, after which the buffers were carefully removed. Three wash steps were carried out, each of 50 μl of ABC buffer and 50 μl acetonitrile and incubation at room temperature. The gel pieces were dried by the addition of 100 μl of 100% acetonitrile for 2 min, which was then discarded. The proteins in the dried gel pieces were then digested with 0.1 μg trypsin (Promega) in 20 μl ABC with incubation overnight at 37°C. The tryptic digest was stopped with 1 μl of a 50% (v/v) formic acid solution and sonication for 15 min. The samples were filtered after the addition of 10 μl of water before transfer into LCMS vials.

The resulting tryptic digest from each gel slice was injected onto a Dionex Nanomate 3000 (ThermoFisher) nano liquid chromatography (LC) system directly coupled to an Orbitrap Fusion Tribrid Mass Spectrometer. The peptides were desalted for 5 min on an Acclaim PepMap C18 (300 Å, 5 mm x 300 μm) trap column at a flow rate of 10 μL/min with loading solvent, and separated on an Acclaim PepMap C18 (100 Å, 150 mm × 0.075 mm) column at a flow rate of 0.3 μL/min at 35°C. A linear gradient from 5% to 40% solvent B over 60 min was employed followed by a wash and re-equilibration with 40-99% B over 5 min, a 5 min hold at 99% B, return to 5% B over 6 min, and held for 7 min. The solvents used were: (A) 0.1% formic acid, 99.9% water; (B) 0.08% formic acid, 80% acetonitrile, 19.92% water. The nano-LC was directly coupled to the Nanospray Flex Ion source of the Orbitrap Fusion MS. The ion spray voltage was set to 2400 V, the sweep gas was set to 1 Arb and the ion transfer tube temperature was set to 300°C. Data were acquired in data-dependent acquisition mode consisting of a Orbitap-MS survey scan followed by parallel acquisition of a high resolution Orbitrap scan at 120,000 resolution and multiple MS/MS events in the linear ion trap, over a 3 second period. First stage MS analysis was performed in positive ion mode over the mass range of m/z 400-1500 with an AGC target of 4 x 10^5^ and a maximum injection time of 50 ms. Tandem mass spectra were acquired in the ion trap on precursor ions that exceeded an intensity threshold of 1000 counts with charge state 2-7. Spectra were acquired using quadrupole isolation with a 1.6 m/z isolation window and (Higher energy Collisional Dissociation) HCD set at 28% based on the size and charge of the precursor ion for optimum peptide fragmentation. Ion trap scan rate was set to rapid with an AGC target of 4 x 10^3^ and a maximum injection time of 300 ms, the instrument was set to utilise the maximum parallelizable time for injecting ions into the trap during a 3 second window whilst the orbitrap was collecting high resolution MS spectra. Dynamic exclusion was set to exclude precursor ions after one occurrence with a 15 second interval and a mass tolerance of 10 ppm.

Analysis of the data for protein identification was conducted using the Sequest algorithm in Proteome Discoverer v2.2 (ThermoFisher). Carbamidomethyl was selected as the alkylating agent and trypsin was selected as the digestion enzyme. Dynamic modifications were selected for oxidation on NifD with a maximum of three modifications. Tandem mass spectrometry data were searched against a database of tryptic peptides for NifD derived from the polyprotein amino acid sequence encoded by SN14 and the *N. benthamiana* proteome, common contaminants and organism specific databases annotated from UniProt. The database search results were curated to yield the protein identifications using a 1% global false discovery rate (FDR) determined by the in-built FDR tool within Proteome Discoverer software.

---

**Supplementary Table 1.**
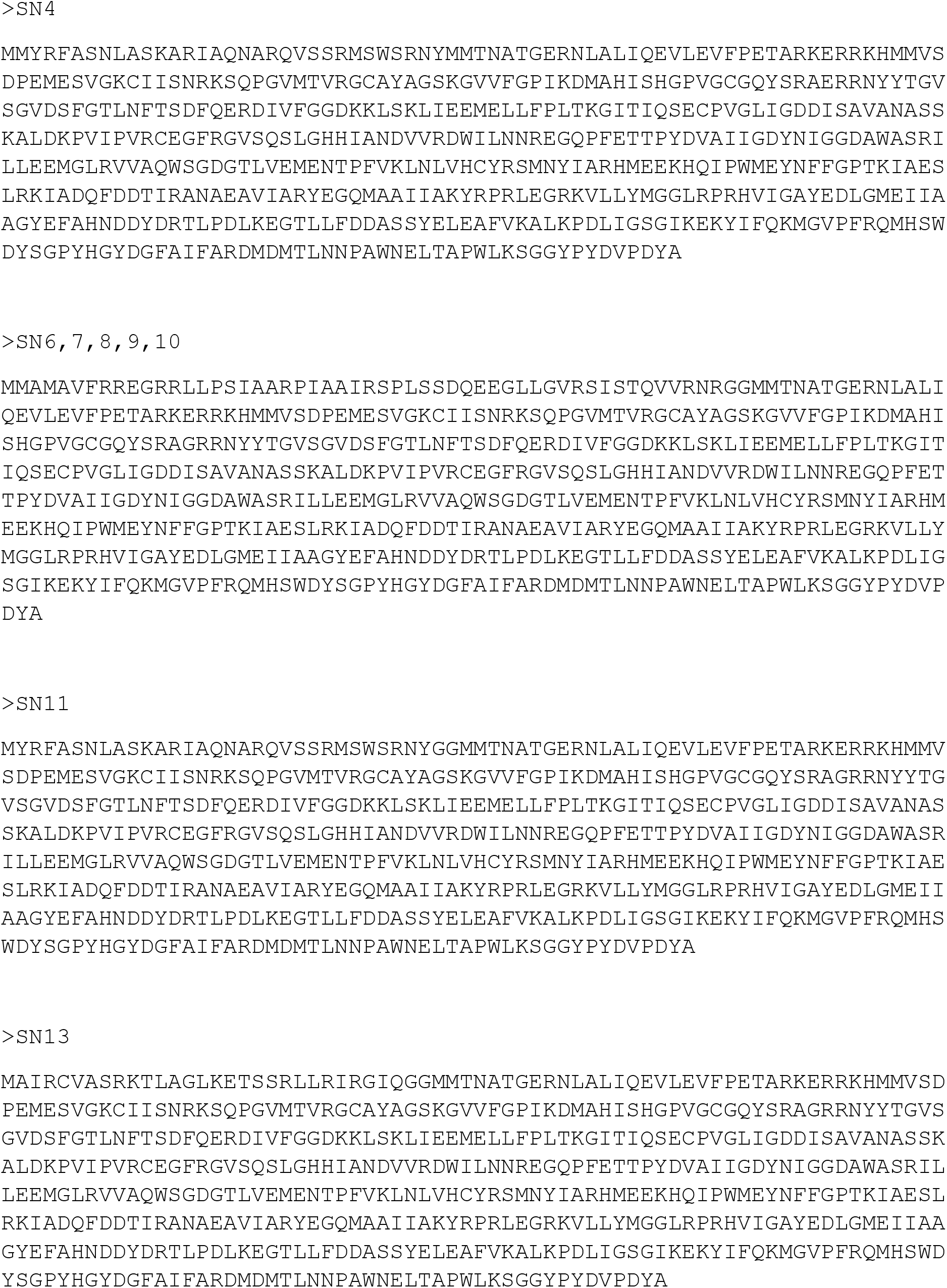
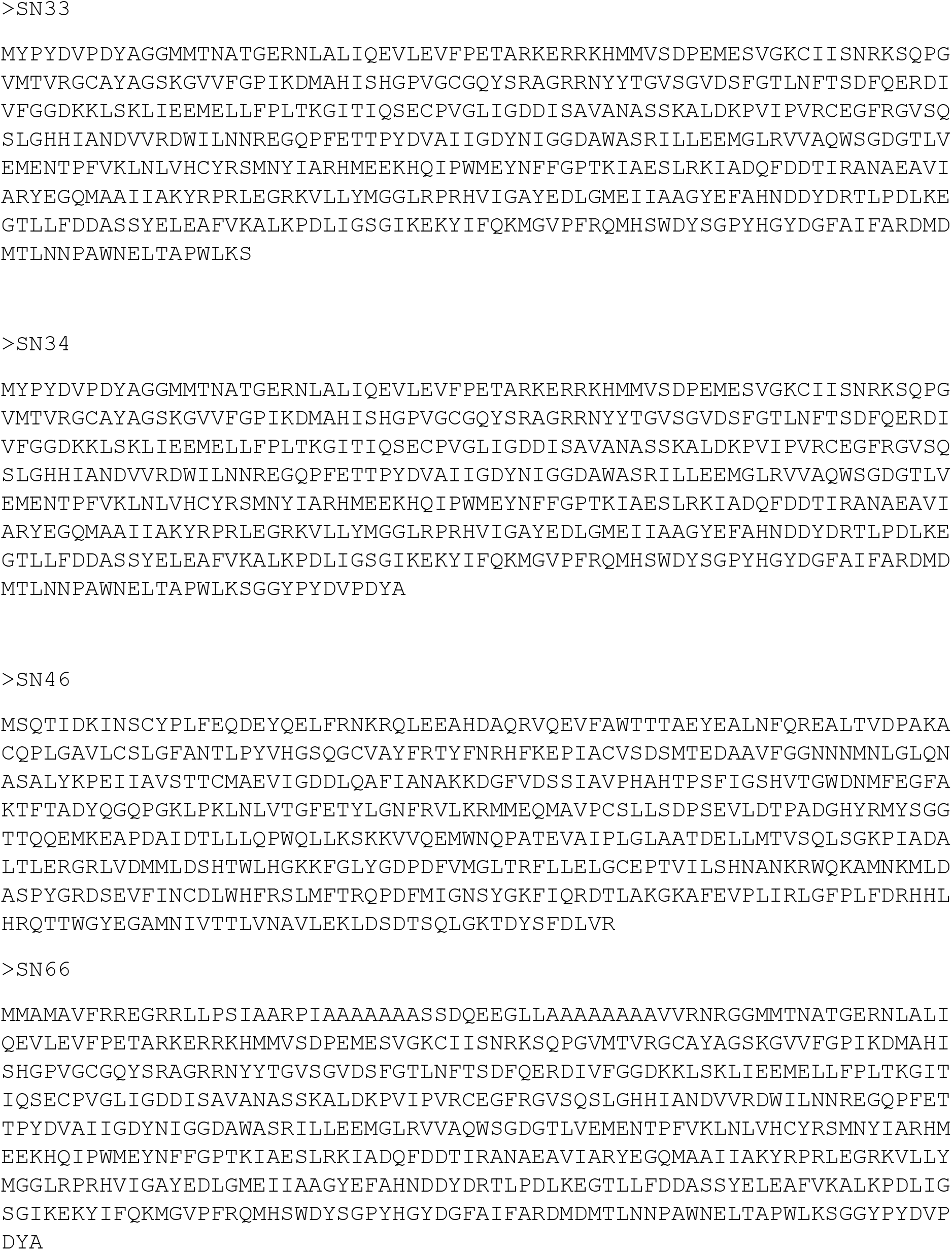
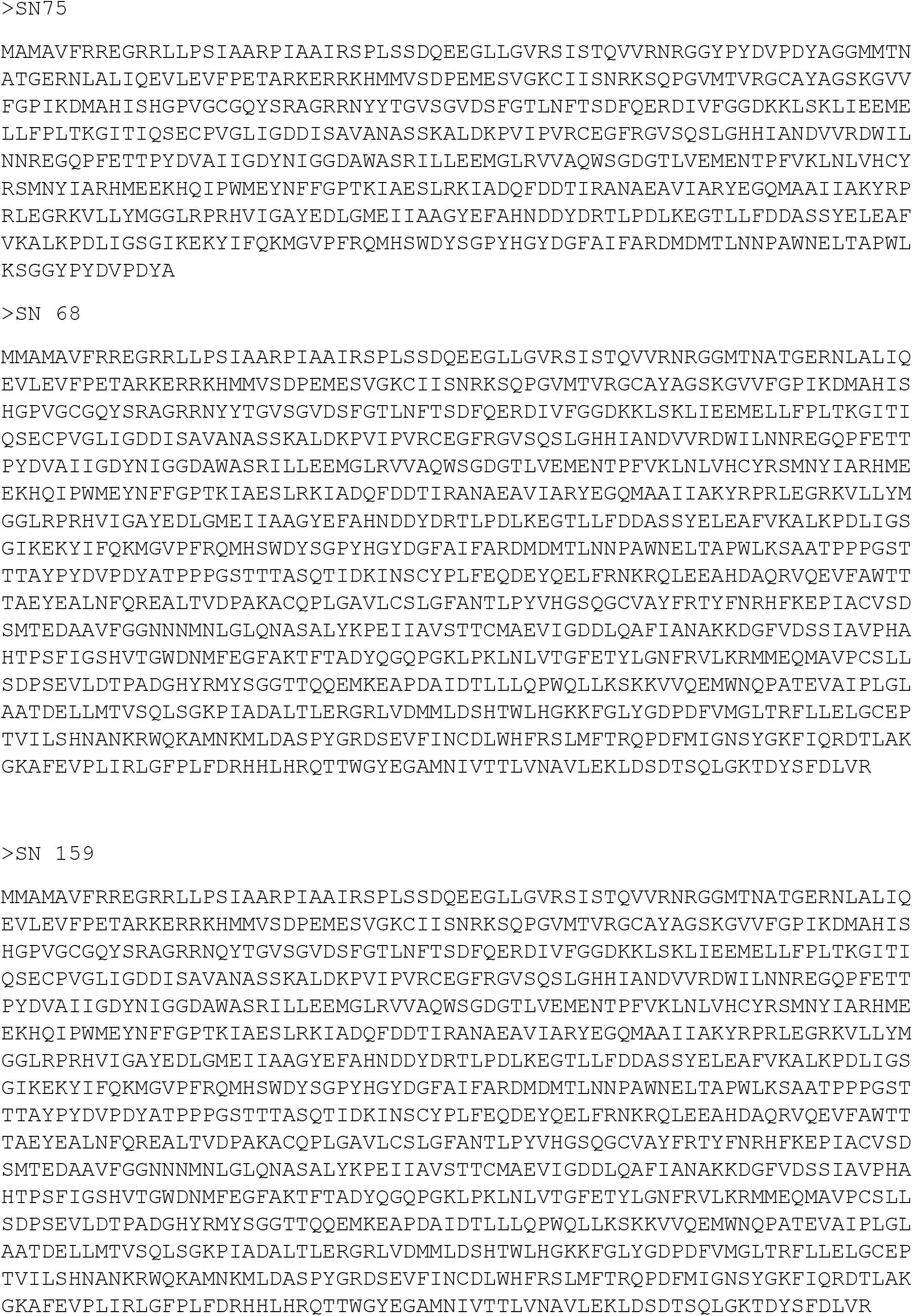
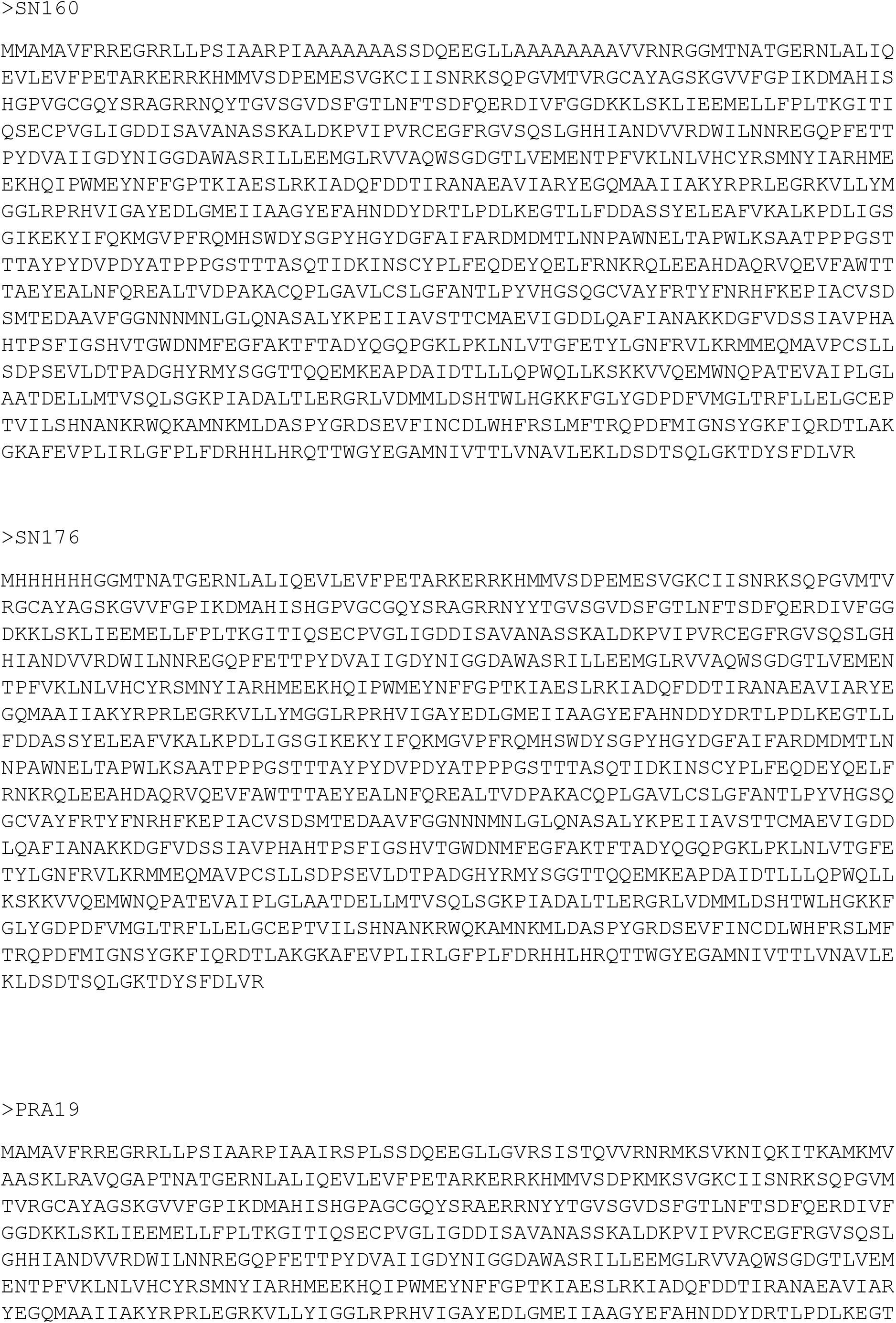
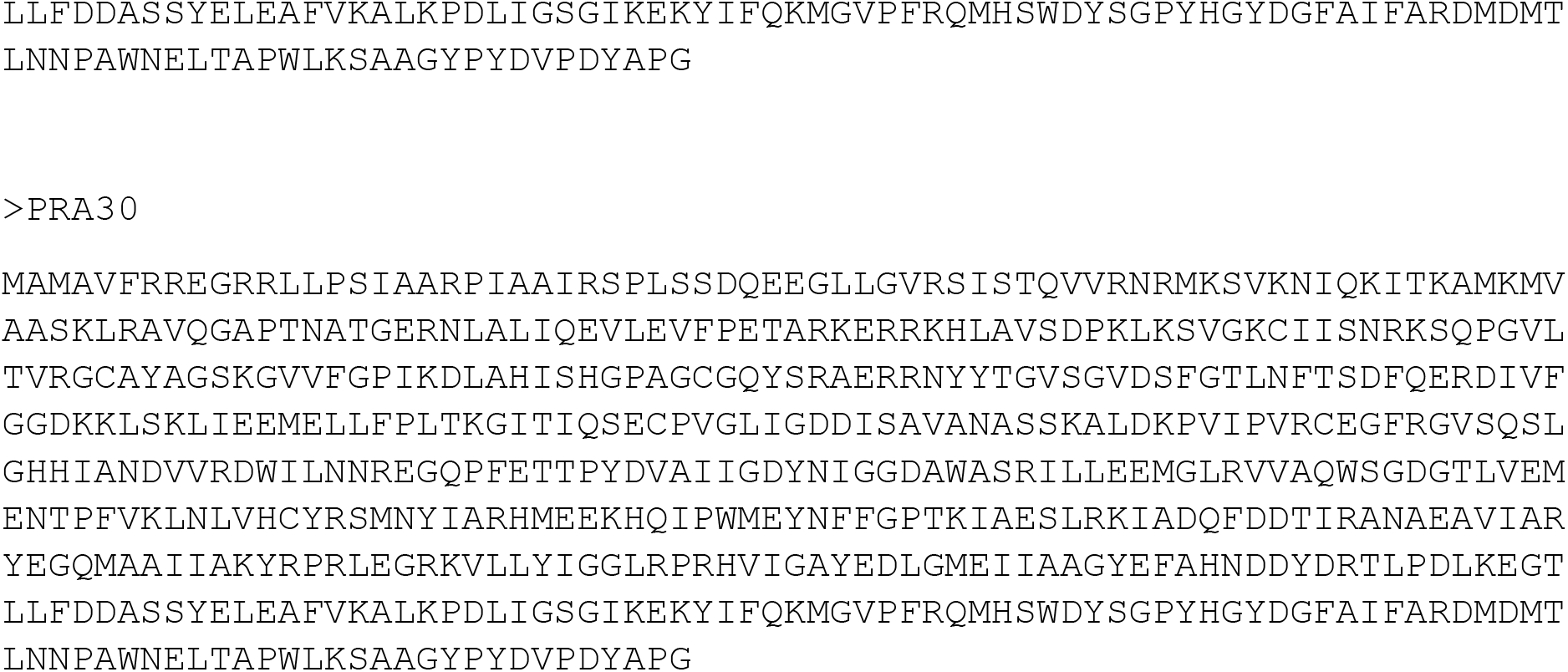
Protein sequences for Nif constructs used in this Study

**Supplementary Table 2.** >Frequency distribution of amino acid residues in the 1476 naturally occurring NifD sequences around the secondary cleavage site. * “X” meaning an unknown amino acid is present in the sequences *Methylocella palustris* (Q6KCQ3) and *Methylosinus trichosporium* (Q6KCQ2).

| *K. oxytoca* amino acid | R97 | R98 | N99 | Y100 | Y101 | T102 |
| --- | --- | --- | --- | --- | --- | --- |
| Position relative to cleavage site | -3 | -2 | -1 | 1 | 2 | 3 |
| most common | R, 1474 (99.86%) | R, 1472 (99.73%) | N, 1444 (97.83%) | Y, 1056 (71.54%) | Y, 951 (64.43%) | V, 402 (27.24%) |
| 2nd most common | Gap, 2 (0.14%) | Gap, 2 (0.14%) | H, 17 (1.15%) | Q, 159 (10.77%) | A, 240 (16.26%) | I, 374 (25.34%) |
| 3rd most common |  | C, 1 (0.07%) | F, 7 (0.47%) | L, 114 (7.72 %) | T, 148 (10.03%) | T, 147 (9.97%) |
| 4th most common |  | G, 1 (0.07%) | A, 3 (0.20%) | K, 108 (7.32%) | M, 44 (2.98%) | K, 135 (9.15%) |
| 5th most common |  |  | X, 2 (0.14%) * | F, 27 (1.83%) | F, 38 (2.57%) | R, 113 (7.66%) |
| 6th most common |  |  | Gap, 2 (0.14%) | M, 10 (0.68%) | G, 27 (1.83%) | N, 96 (6.50%) |
| 7th most common |  |  | S, 1 (0.07%) | E, 1 (0.07%) | S, 14 (0.95%) | D, 68 (4.61%) |
| 8th most common |  |  |  | Gap, 1 (0.07%) | V, 12 (0.81%) | S 40 (2.71%) |
| 9th most common |  |  |  |  | N, 1 (0.07%) | E, 27 (1.83%) |
| 10th most common |  |  |  |  | Gap, 1 (0.07%) | Q, 23 (1.56%) |
| 11th most common |  |  |  |  |  | L, 20 (1.36%) |
| 12th most common |  |  |  |  |  | A, 12 (0.81%) |
| 13th most common |  |  |  |  |  | H, 9 (0.81%) |
| 14th most common |  |  |  |  |  | M, 8 (0.61%) |
| 15th most common |  |  |  |  |  | Y, 2 (0.14%) |

